# FSTruct: an *F*_*ST*_ -based tool for measuring ancestry variation in inference of population structure

**DOI:** 10.1101/2021.09.24.461741

**Authors:** Maike L. Morrison, Nicolas Alcala, Noah A. Rosenberg

## Abstract

In model-based inference of population structure from individual-level genetic data, individuals are assigned membership coefficients in a series of statistical clusters generated by clustering algorithms. Distinct patterns of variability in membership coefficients can be produced for different groups of individuals, for example, representing different predefined populations, sampling sites, or time periods. Such variability can be difficult to capture in a single numerical value; membership coefficient vectors are multivariate and potentially incommensurable across groups, as the number of clusters over which individuals are distributed can vary among groups of interest. Further, two groups might share few clusters in common, so that membership coefficient vectors are concentrated on different clusters. We introduce a method for measuring the variability of membership coefficients of individuals in a predefined group, making use of an analogy between variability across individuals in membership coefficient vectors and variation across populations in allele frequency vectors. We show that in a model in which membership coefficient vectors in a population follow a Dirichlet distribution, the measure increases linearly with a parameter describing the variance of a specified component of the membership vector. We apply the approach, which makes use of a normalized *F*_*ST*_ statistic, to data on inferred population structure in three example scenarios. We also introduce a bootstrap test for equivalence of two or more groups in their level of membership coefficient variability. Our methods are implemented in the R package FSTruct.

## Introduction

In the past two decades, computational methods for inference of population structure from individual-level genetic data have contributed a rich and informative set of approaches for the analysis of genetic variation. Model-based clustering methods such as Admixture (Alexander *et al*., 2009; Alexander and Lange, 2011), Baps (Corander *et al*., 2004, 2008) and Structure (Falush *et al*., 2003, 2007; Hubisz *et al*., 2009; Pritchard *et al*., 2000) are now routinely used to generate insights into population structure and evolutionary history in diverse species of interest in ecology, evolution, conservation biology, and agriculture (Guillot and Orlando, 2017).

In model-based inference of population structure, individuals are clustered based on their multilocus genotypes into a series of statistical clusters, such that each individual possesses a membership coefficient for each cluster. Each membership coefficient represents the proportion of an individual’s ancestry that is derived from the associated cluster. Interpreting the membership coefficients of individuals from various predefined populations, sampling sites, or other groups of biological interest can illuminate patterns of genetic variation and population structure. Researchers often investigate variability of membership patterns within predefined groups, as well as similarities and differences in the membership patterns of distinct groups.

One type of comparison that is frequently of interest is an assessment of relative levels of variation in membership coefficients among the individuals belonging to two or more predefined groups. This type of comparison arises in many contexts, such as when exploring differences in membership variability between admixed and non-admixed populations, between populations from different time periods, or between different types of data on the same sampled individuals.

For example, in a study of ancient human DNA samples dating over a period of hundreds to thousands of years, Antonio *et al*. (2019) sought to examine if the population of Rome possessed greater diversity in ancestry during certain periods of the Roman Empire. They estimated membership coefficients using Admixture and interpreted the inferred coefficients to claim that during the Imperial Rome period, when the Roman Empire was at its peak, ancestry was more variable than during earlier periods, when Rome was more isolated (Antonio *et al*., 2019, Fig. 1).

**Figure 1:**
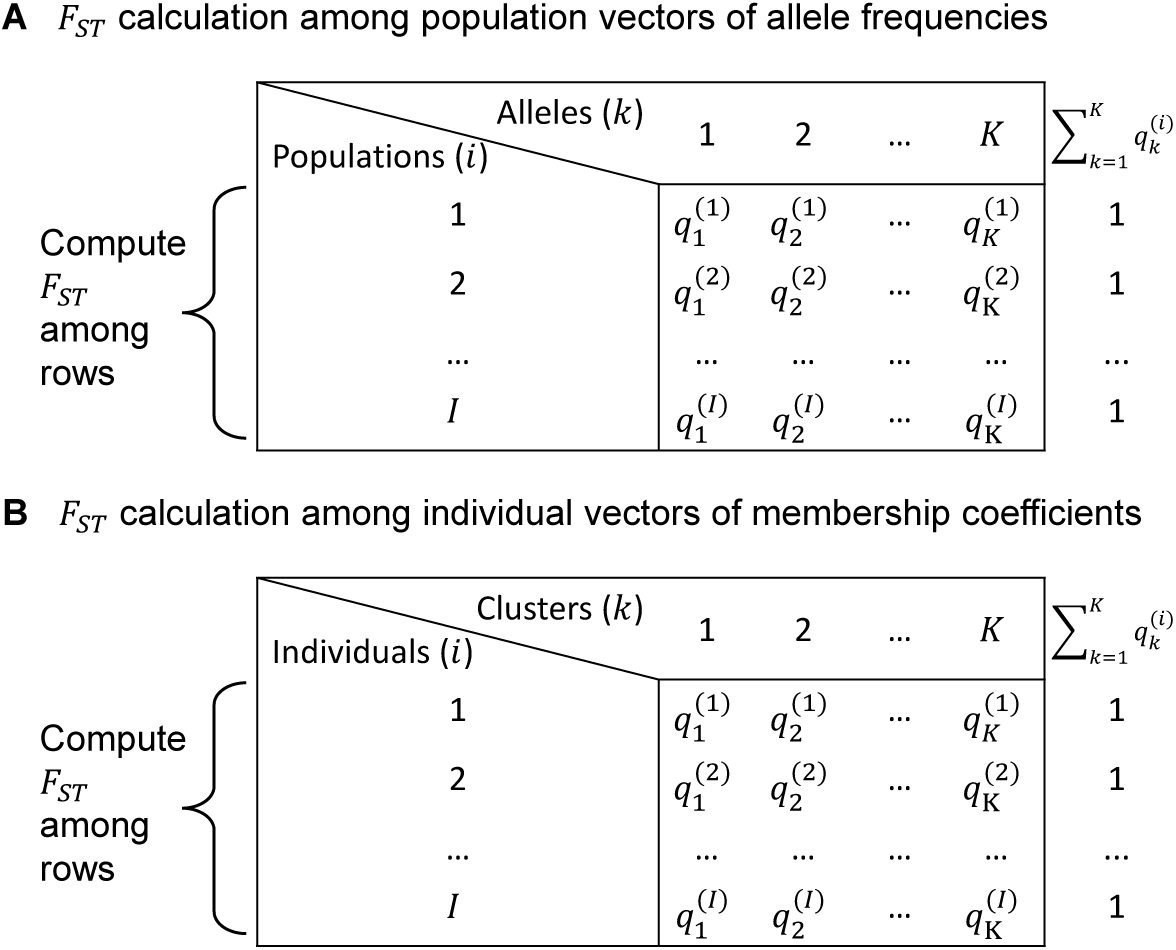
The analogy of the use of *F*_*ST*_ to measure membership variability. **(A)** A standard application of *F*_*ST*_ to measure variability of allele frequency vectors across populations; 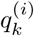 is the frequency of allele *k* in population *i*. **(B)** Use of *F*_*ST*_ to measure variability of membership coefficient vectors across individuals; 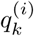 is the membership coefficient of individual *i* in cluster *k*. The matrix containing entries 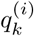 is a *Q* matrix.

Interpretations of inferred membership coefficients to make relative claims about membership variability have generally relied on visual assessment of population structure diagrams rather than on statistical hypothesis testing. In particular, as in Antonio *et al*. (2019), researchers seeking to quantify variability in membership coefficients across individuals or to compare this variability between two or more groups often do so visually or informally.

Here, we introduce a statistical method to measure variability in membership coefficients inferred by model-based clustering, and to compare this variability across populations. We apply the method to examples from real and simulated data. The method is implemented in the *R* package FSTruct.

## Materials & Methods

### Overview

The output of population structure inference software programs such as Structure and Admixture is a representation of individual membership coefficients in matrix form. The matrix, often denoted *Q* and termed a “*Q* matrix,” has *I* rows, corresponding to *I* individuals, and *K* columns, corresponding to the total number of clusters (Figure 1B). The entry in row *i* and column *k*, 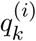, represents the membership coefficient of individual *i* in cluster *k*: the proportion of the ancestry of individual *i* that is assigned to cluster *k*. Each row sums to 1, or 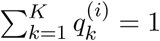 for each *i*.

We seek to compute a measure of variability among ancestry vectors for individuals, that is, among rows of *Q*. We wish for the measure to be comparable across different data sets, possibly representing different samples. This problem is complicated by the fact that different *Q* matrices might include different numbers of clusters; furthermore, column entries for some clusters might vary greatly across individuals while other columns are more uniform.

We approach the problem by modifying the population differentiation statistic *F*_*ST*_ to fit this ancestry scenario. *F*_*ST*_ measures allele frequency variability among subpopulations, and it is computed using a set of allele frequency vectors that each sum to 1. This setting is mathematically analogous to *Q* matrices, in which vectors of membership coefficients for each individual sum to 1. In the analogy, each individual represents a “population,” and its cluster membership is analogous to an “allele frequency” (Figure 1).

By computing *F*_*ST*_ among individual vectors of membership coefficients, we can measure the variability of a single *Q* matrix. To facilitate comparisons of *Q* matrices with different numbers of individuals or clusters, we use a normalization of *F*_*ST*_. Despite the general understanding that *F*_*ST*_ can in principle reach 1, features of a data set constrain the maximal value of *F*_*ST*_, so that the maximum is often less than 1 (Jakobsson *et al*., 2013; Alcala and Rosenberg, 2017, 2019). The constrained maximum is relatively low when *I*, the number of individuals in a *Q* matrix, is small (analogous to a small number of populations), or when *M*, the mean membership of the highest-membership ancestry cluster, is close to 0 or 1 (analogous to an extreme value for the frequency of the most frequent allele). Denoting this maximum 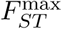, we normalize *F*_*ST*_ by its maximum, using the ratio 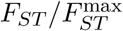 as a measure of variability that is comparable across *Q* matrices of different size. This measure ranges between 0 and 1, equaling 0 when members of a population have identical ancestry and equaling 1 when vectors of membership coefficients are maximally variable.

### The 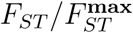 formula

Consider a scenario with *I* subpopulations and *K* distinct alleles. Allele *k* has frequency 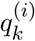 in subpopulation *i*, with 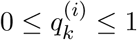 and 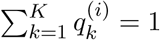.

To calculate *F*_*ST*_ among the *I* subpopulations, we use *F*_*ST*_ = (*H*_*T*_ − *H*_*S*_)*/H*_*T*_, where *H*_*S*_ represents the mean heterozygosity of the subpopulations and *H*_*T*_ represents the heterozygosity of the total population formed by pooling the subpopulations.

The subpopulation heterozygosity *H*_*S*_ is the mean expected frequency of heterozygotes across all *I* subpopulations, or 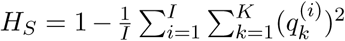. The total heterozygosity *H*_*T*_ is the expected frequency of heterozygotes under Hardy-Weinberg equilibrium in a population whose allele frequencies equal the mean allele frequencies across subpopulations: 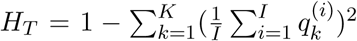. The quantity 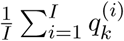 gives the mean frequency of allele *k* across subpopulations.

With the total population assumed to be polymorphic so that *H*_*T*_ > 0, for the setting of *I* subpopulations and *K* alleles, with *K* possibly arbitrarily large, Alcala and Rosenberg (2021) obtained the maximal value possible for *F*_*ST*_ given a fixed value of 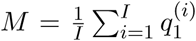, where allele *k* = 1 represents the allele of greatest mean frequency across the *I* subpopulations. Writing 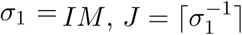, and {*σ*_1_} = *σ*_1_ − ⌊*σ*_1_⌋, we have (Alcala and Rosenberg, 2021, eq. 3)

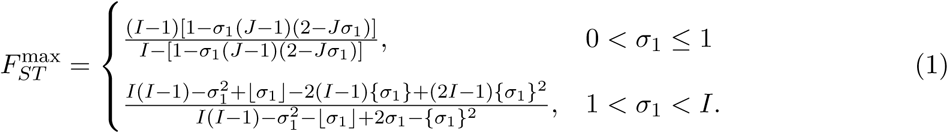

This maximum is plotted as a function of *M* for five different values of *I* in Figure 2.

**Figure 2:**
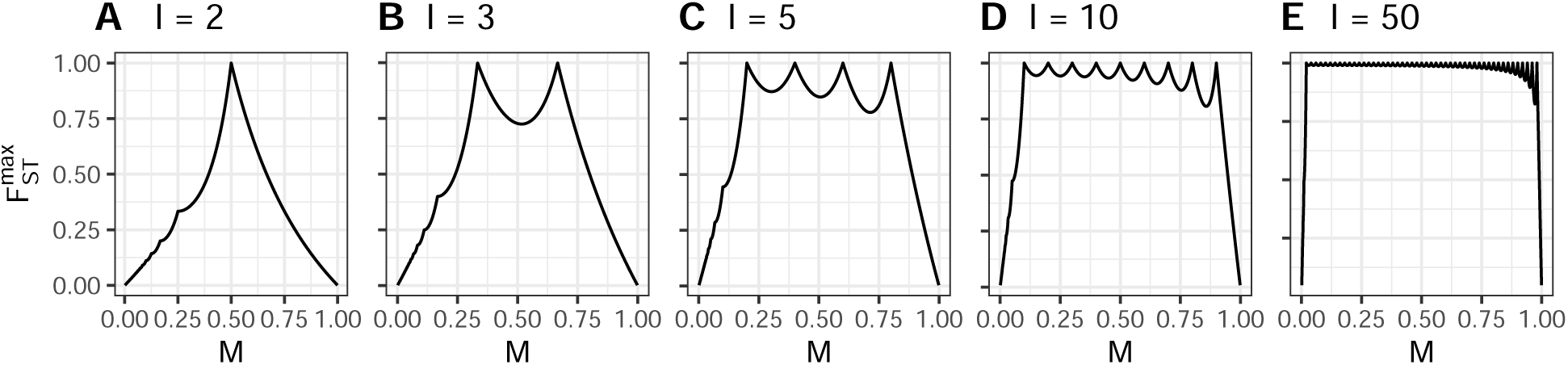
Bounds on *F*_*ST*_ as a function of *M*, the frequency of the most frequent allele—or the ancestry cluster of greatest membership, in our analogy. Bounds are evaluated using eq. 1 for different values of *I*, the number of populations (or the number of individuals, in our analogy). (A) *I* = 2. (B) *I* = 3. (C) *I* = 5. (D) *I* = 10. (E) *I* = 50.

In the language of our analogy, *I* is the number of individuals—the number of rows in the *Q* matrix; *M* is the sample mean membership coefficient for the most frequent ancestral cluster across all *I* individuals; and *σ*_1_ = *IM* is the largest entry in the vector that sums column entries of the *Q* matrix across rows. The latter case of eq. 1, with 1 < *σ*_1_ < *I*, is generally more relevant in the setting of population clustering, as *I* is typically larger than *K*, so that 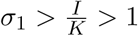.

The ratio 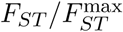, which represents a normalized measure of variability that can be compared among different groups of individuals with different values of *I* or *K* or both, ranges between 0 and 1, taking a value of 0 when all individuals in a group have identical membership coefficients. It has a value of 1 when they are as variable as possible given *M*.

Alcala and Rosenberg (2021) showed that for 0 < *σ*_1_ ≤ 1, the maximum is realized when each ancestry cluster is found in only a single individual and each individual has exactly *J* ancestry clusters with coefficients greater than zero: *J* − 1 clusters with coefficients of *M*, one cluster with a coefficient of 1 − (*J* − 1)*σ*_1_ ≤ *M*, and all others with coefficients of 0.

For 1 < *σ*_1_ < *I*, the maximum is realized when only the ancestry cluster of greatest membership is shared among individuals, and at most a single individual contains ancestry from multiple sources. More formally, this scenario occurs when ⌊*σ*_1_⌋ individuals possess all of their membership in the cluster of greatest membership, a single individual has membership coefficient {*σ*_1_} for the cluster of greatest membership and coefficient 1 −{*σ*_1_} for one other cluster, and the remaining *I* −⌊*σ*_1_⌋− 1 individuals each have membership coefficient 1 for mutually distinct ancestry clusters.

### Statistical test to compare values of 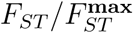

In applications, we may wish not only to compute 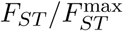 for a single population, but also to compare this ratio between two or more populations using a statistical test. We accomplish this task by bootstrap resampling of rows to generate replicate *Q* matrices for each population. We then compute the 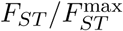 statistic for each of these replicate matrices. This process generates a bootstrap distribution of the statistic for each population. We then use a Wilcoxon rank-sum test to determine if pairs of bootstrap distributions of the statistic for different sets of individuals are significantly different; we use a Kruskal-Wallis test to compare three or more sets of individuals.

### Software availability

We have implemented the method in the R package FSTruct (pronounced “F-struct”), which is available for download from github.com/MaikeMorrison/FSTruct. This package includes functions that compute 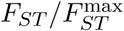 from a *Q* matrix such as those produced by Admixture or Structure, generate bootstrap replicates and distributions for arbitrarily many *Q* matrices, and visualize *Q* matrices.

## Results

### Simulation examples

#### Dirichlet model

To illustrate our method, we used individual membership coefficient vectors drawn from a Dirichlet distribution (Kotz *et al*., 2000). This distribution is suited for use as the underlying model for finite vectors of nonnegative numbers (*q*_1_, *q*_2_, …, *q*_*K*_) that sum to one, 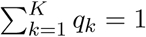, and it has appeared in previous studies of membership coefficient vectors (Pritchard *et al*., 2000; Huelsenbeck and Andolfatto, 2007).

We treat individual membership coefficient vectors in a population as following a Dirichlet distribution with parameter vector *α****λ*** = *α*(*λ*_1_, *λ*_2_, …, *λ*_*K*_), where 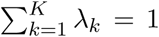. We denote this distribution by Dir(*α*(*λ*_1_, *λ*_2_, …, *λ*_*K*_)). Here, ***λ*** is a vector of length *K* whose elements determine the parametric mean membership coefficient for each ancestral cluster. The value of *α* controls the variance of *q*_*k*_, the individual membership coefficient in cluster *k*: Var[*q*_*k*_] = *λ*_*k*_(1 − *λ*_*k*_)*/*(*α* + 1). Thus, an increase in *α* lowers the variances of membership coefficients.

To generate a random *Q* matrix with *I* individuals and *K* ancestry clusters, we draw *I* independent and identically distributed Dir(*α*(*λ*_1_, *λ*_2_, …, *λ*_*K*_)) vectors, (*q*_1_, …, *q*_*K*_), which each comprise a set of membership coefficients for a single individual. Each vector is a row of the simulated *Q* matrix and is a draw from a Dirichlet distribution with mean membership coefficients (*λ*_1_, *λ*_2_, …, *λ*_*K*_). Variability of membership coefficients across individuals is controlled by *α*. Hence, we proceed by (1) using the Dirichlet distribution to simulate *Q* matrices with specified membership coefficient means and variances, (2) computing 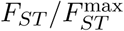 for each *Q* matrix, and (3) examining the relationship between the value of 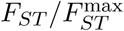 for each *Q* matrix and the variance of the Dirichlet distribution used to simulate it.

#### Dirichlet simulations

To investigate the behavior of 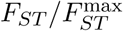 in relation to a measure of variability in membership coefficients, we used the Dirichlet distribution to simulate *Q* matrices with known variability. We simulated *Q* matrices with *I* = 50 individuals and *K* = 2 clusters. Each simulation replicate thus drew *I* = 50 ancestry vectors from a Dir(*α*(*λ*_1_, *λ*_2_)) distribution.

We fixed 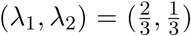, so that membership in cluster 1 has mean 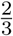 across individuals in a population and membership in cluster 2 has mean 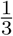. The variance of the membership coefficient for a specific cluster, across sampled individuals, then equals 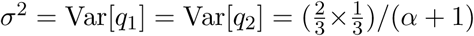; both coefficients have the same variance. As *α* ranges in (0, ∞), the variance ranges in 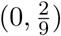.

We performed 500 replicate simulations of samples of 50 individuals for each of 45 values of *α*, choosing *α* values to obtain variances 0.001, 0.005, 0.01, 0.015, …, 0.22, ranging from near the lower bound of 0 on the variance and stopping short of the upper bound of 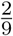.

Next, we compared the value of 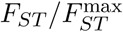 for each simulated *Q* matrix to the variance parameter of the Dirichlet distribution used to generate it. As 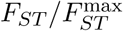 measures variability of *Q* matrices, we expect to see a positive relationship between the Dirichlet variance used to generate the *Q* matrix and our estimate of its variability,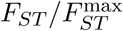.

Simulation results, depicting the 500 values of 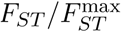 for each of the 45 choices of the Dirichlet variance 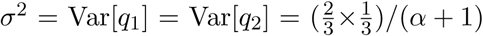, appear in Figure 3. In the figure, the relationship between 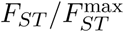 and *σ*^2^ is strongly linear, with slope 4.5.

**Figure 3:**
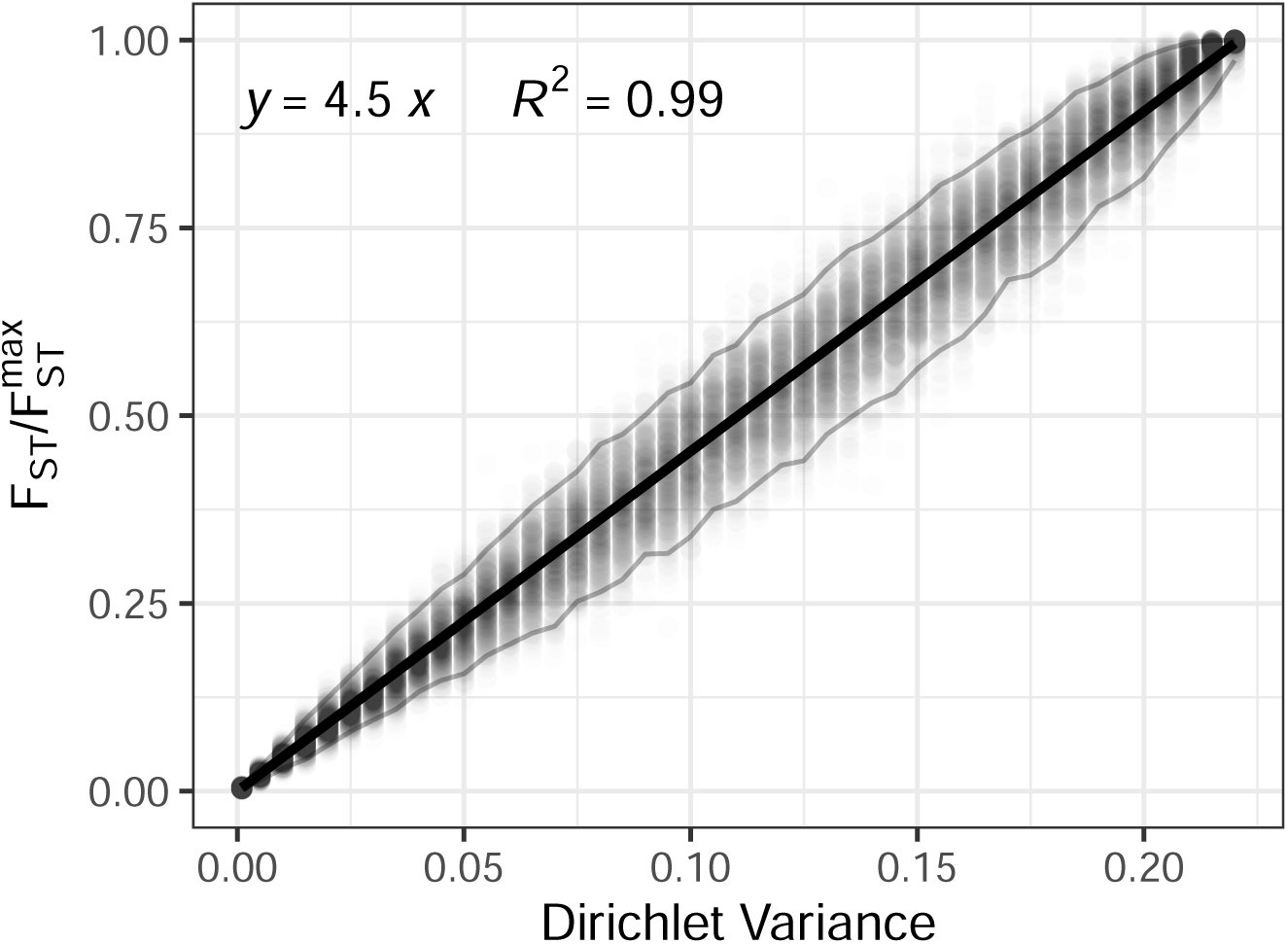
Linear relationship between 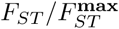 and Var[*q*_1_] = Var[*q*_2_], the variance across individuals of individual membership coefficients under a Dirichlet distribution. For each of 45 values of Var[*q*_1_] = Var[*q*_2_], 500 points are plotted, each representing a random *Q* matrix with dimensions 50 × 2. Rows of the *Q* matrix are simulated using a Dirichlet distribution with means 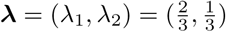 and variances Var[*q*_1_] = Var[*q*_2_] = *λ*_1_*λ*_2_*/*(*α* + 1), with *α* chosen to produce variances 0.001, 0.005, 0.01, 0.015, … 0.22. Each *Q* matrix gives rise to an associated value of 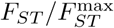, plotted on the vertical axis. A regression line fit to the 500 × 45 points with intercept 0 has slope 4.5, or 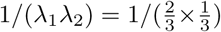, and it explains 99% of the variability in 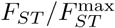. Grey lines mark the 2.5% and 97.5% percentiles, and thus contain 95% of the points.

Noticing that the empirical slope, 4.5, was the reciprocal of 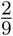, the upper bound of the Dirichlet variance, we sought to obtain a mathematical relationship between 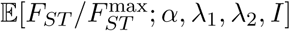, the expectation of 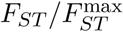 under the Dirichlet model, and the parametric variance of each membership coefficient in the model. This calculation, performed in the Appendix, confirms the relationship (eq. A.11)

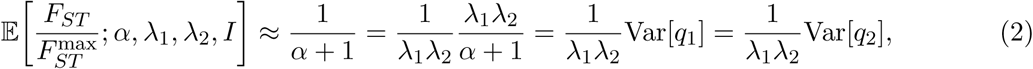

where 1*/*(*λ*_1_*λ*_2_) = 4.5 in the example plotted in Figure 3. Thus, the simulations and an analytical calculation confirm that in a simple Dirichlet model, the 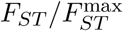 measure has a linear relationship with the variance across sampled individuals of membership coefficient *q*_1_ (or *q*_2_). Importantly, the expected value of 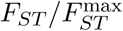 in eq. 2 is independent of the mean membership coefficients, depending only on the Dirichlet parameter *α*, which controls variability. This result supports the use of 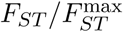 to measure variability in populations that possess different mean membership coefficients.

#### Visual illustration of values of 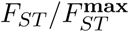

Continuing with the Dirichlet simulations, we next sought to visually illustrate the relationship of 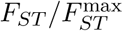 to the variance and mean of membership coefficients. We considered *Q* matrices with four different values of *α*, representing four levels of variance in membership coefficients, and two different vectors for the parametric mean membership coefficients ***λ***. For each of the eight settings (four variances, two means), we considered two *Q* matrices.

These eight simulated pairs of *Q* matrices are visualized in Figure 4A and 4D respectively, where they are colored according to the value of the *α* parameter used to simulate them. For the lowest-variability case (*α*_1_, red), the simulated individual membership coefficients show little deviation from the mean, 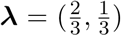 for 4A and 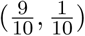 for 4D. As the variance parameter increases (*α*_2_, purple; *α*_3_, blue), variance in membership coefficients is increasingly visible. For the highest-variability case (*α*_4_, green), membership coefficients are centered on (*λ*_1_, *λ*_2_) = (1, 0) for approximately 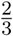 or 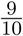 of the individuals, and on (*λ*_1_, *λ*_2_) = (0, 1) for the remaining individuals.

**Figure 4:**
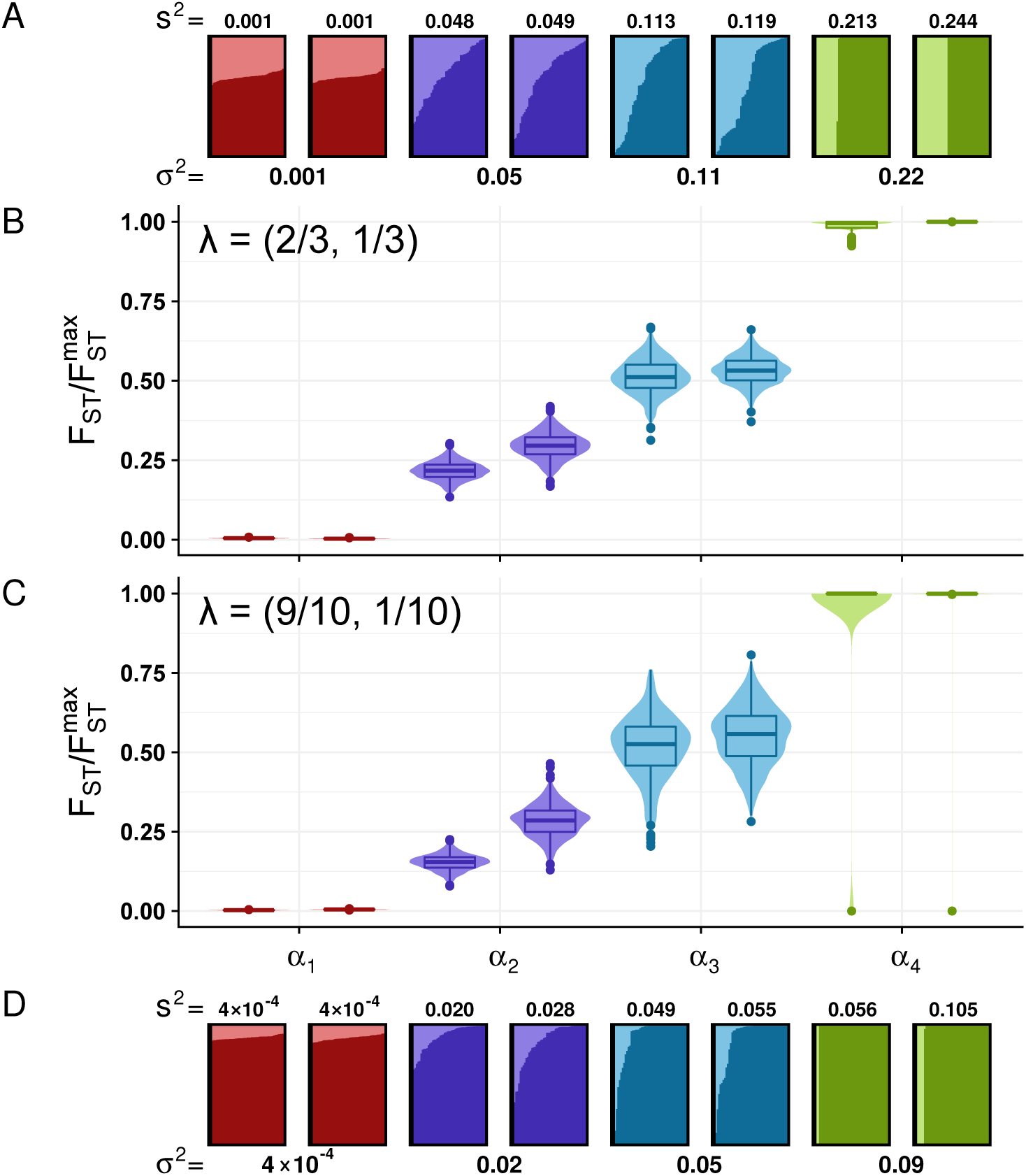
Dependence of bootstrap distributions of 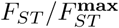 for simulated *Q* matrices on the Dirichlet variance parameter *α*, rather than the Dirichlet mean *λ*. **(A, D)** *Q* matrices simulated using specified Dir(*α****λ***) distributions. **(B, C)** Bootstrap distributions of 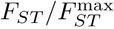 for *Q* matrices from (A) and (D), plotted directly below or above the corresponding matrix. In both (A) and (D), eight matrices were simulated, two for each of four values of *α* selected to span the range of variances: *α*_1_ = 21901*/*99, *α*_2_ = 341*/*99, *α*_3_ = 101*/*99, *α*_4_ = 1*/*99. Matrices are annotated by associated variances *σ*^2^ = *λ*_1_*λ*_2_*/*(*α* + 1). In (A), matrices are simulated with parametric mean 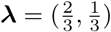 and are taken from matrices plotted in Figure 3. In (D), matrices are simulated with a more extreme parametric mean, 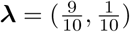. Each vertical bar represents an individual membership coefficient vector (*q*_1_, *q*_2_); the proportion of each bar colored a darker shade represents *q*_1_ and the proportion in a lighter shade corresponds to *q*_2_. The parametric variance of a *Q* matrix, *σ*^2^ = *λ*_1_*λ*_2_*/*(*α* + 1), ranges in 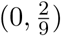 for 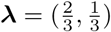 and in (0, 0.09) for 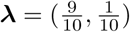. The empirical variance s^2^ is computed for each matrix using the sample mean 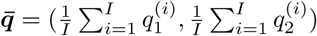 in place of the parametric mean ***λ***. The values of 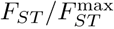 for the eight matrices in (A) are 0.004 and 0.005 for the two simulated with *α*_1_, 0.203 and 0.230 for *α*_2_, 0.496 and 0.461 for *α*_3_, and 1.000 and 0.997 for *α*_4_. The values of 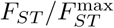 for the eight matrices in (D) are 0.003 and 0.005 for *α*_1_, 0.157 and 0.287 for *α*_2_, 0.539 and 0.571 for *α*_3_, and 1.000 and 1.000 for *α*_4_. In (B) and (C), each bootstrap distribution includes 1,000 bootstrap samples of the *I* = 50 individuals in the associated *Q* matrix.

Bootstrap distributions of 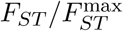 appear in Figure 4B for 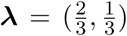 and in 4C for 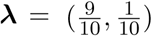. In these panels, we observe that 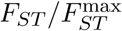 increases from the lowest-variability case (*α*_1_) to the highest-variability case (*α*_4_), in accord with the interpretation that 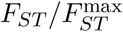 measures variability in membership coefficients.

Comparing panel 4B with panel 4C, we observe that the value of 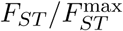 is similar between matrices simulated with the same Dirichlet *α* parameter, irrespective of the mean membership coefficient vectors (***λ***) used to simulate the matrices. This pattern accords with the interpretation that 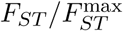 is driven by the variance of membership coefficients and not the mean—as reflected in the analytical result in eq. 2 that under the Dirichlet model, 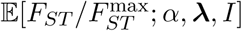 can be written so that it depends on *α* but not on ***λ***.

We also observe in Figure 4A and 4D that the variability among *Q* matrices simulated from the Dirichlet distribution with identical parameters—as reflected in comparisons of pairs of matrices of the same color within a panel—increases with *α*. This variability can lead *Q* matrices simulated with the same parameters to possess quite different sample means and variances, as is the case particularly for the two pairs of matrices simulated with *α*_4_ in Figure 4D. Despite this sampling variability of *Q* matrices under the Dirichlet model, we observe that 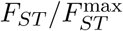, which is largely driven by the underlying parameter *α*, is relatively stable across pairs of *Q* matrices.

### Data examples

To illustrate the application of FSTruct, we apply the method to data examples that represent each of three distinct scenarios in which ancestry variability is of interest: (1) ancestry comparisons of admixed and non-admixed populations, (2) ancestry comparisons of populations representing different time periods or spatial locations, and (3) ancestry comparisons of distinct data sets corresponding to different sets of loci for the same individuals.

#### Admixed populations

A characteristic feature of recently admixed populations is that individuals vary greatly in their ancestry, with some individuals possessing most of their ancestry from one source population, and others possessing most of their ancestry from another source (Verdu and Rosenberg, 2011; Gravel, 2012). Thus, in examining inferred cluster memberships, admixed populations might be expected to give rise to greater variability in ancestry than non-admixed populations.

We therefore evaluated 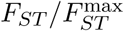 in three populations from an Admixture analysis performed by Verdu *et al*. (2017). The populations include an admixed population from Cape Verde, and Gambian and Iberian populations taken to represent African and European sources for the admixed population. The inferred genetic structure for the three populations is redrawn in Figure 5A.

**Figure 5:**
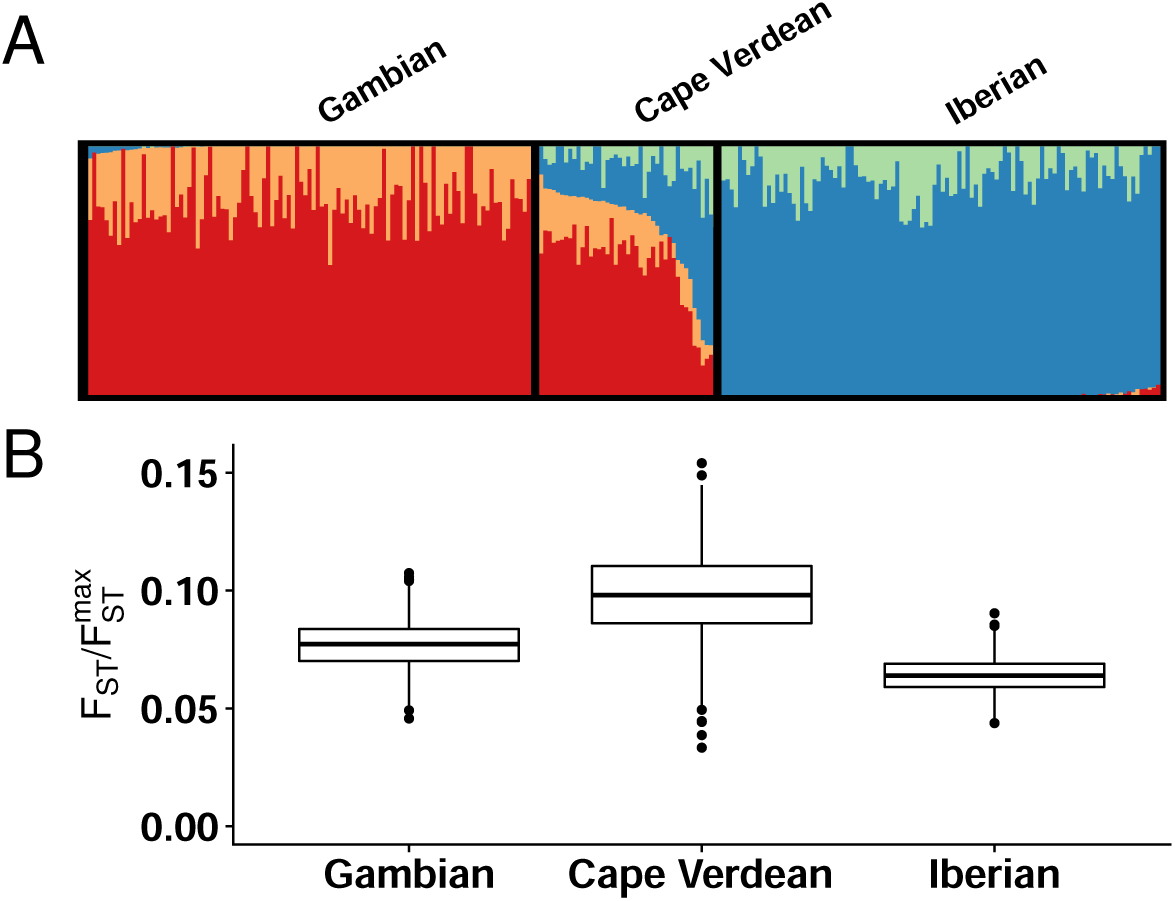
Variability of ancestry in admixed and non-admixed populations. **(A)** *K* = 4 Admixture analysis of Gambian (*n* = 109), Cape Verdean (*n* = 44), and Iberian (*n* = 107) samples. Adapted from Verdu *et al*. (2017). **(B)** Bootstrap distributions of the ancestry variability measure, 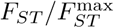, for each population (1,000 replicates).

We computed 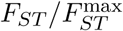 for each of the three populations, measuring ancestry variability of the inferred cluster memberships within each of the three groups. For the non-admixed source populations, this quantity is 0.078 for the Gambian population and 0.064 for the Iberian population (Figure 5B). The value for the admixed Cape Verdean population is greater, equaling 0.100. Pairs of bootstrap distributions of 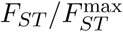 are significantly different (*p* < 2 × 10^−16^ for all three pairwise combinations, Wilcoxon rank sum test). The admixed Cape Verdean population is indeed observed to have greater variability in ancestry according to the 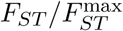 measure than the putative source populations, supporting the use of the measure to distinguish clustering patterns in admixed and non-admixed populations.

#### Populations over time or space

Geographic movements of populations shape patterns of genetic ancestry for samples collected in different spatial locations or from the same location in different time periods. Locations or time periods whose samples contain individuals from many different sources or from recently admixed populations are expected to have highly variable ancestry, whereas locations or periods in which mixing of disparate populations is less salient are expected to have more homogeneous ancestry.

To explore an example of ancestry variability over time, we evaluated 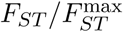 in a Structure analysis conducted by Antonio *et al*. (2019) on samples from 29 archaeological sites near Rome spanning the last 12,000 years. These samples represent eight time periods: Mesolithic, Neolithic, Copper Age, Iron Age and Roman Republic, Imperial Rome, Late Antiquity, Medieval and Early Modern, and the present. The plot of the inferred genetic structure for these samples is redrawn in Figure 6A. Antonio *et al*. (2019) argued, based in part on their version of Figure 6A, that ancestry was variable during the Iron Age and Roman Republic, and highly variable during the Imperial Rome and Late Antiquity periods.

**Figure 6:**
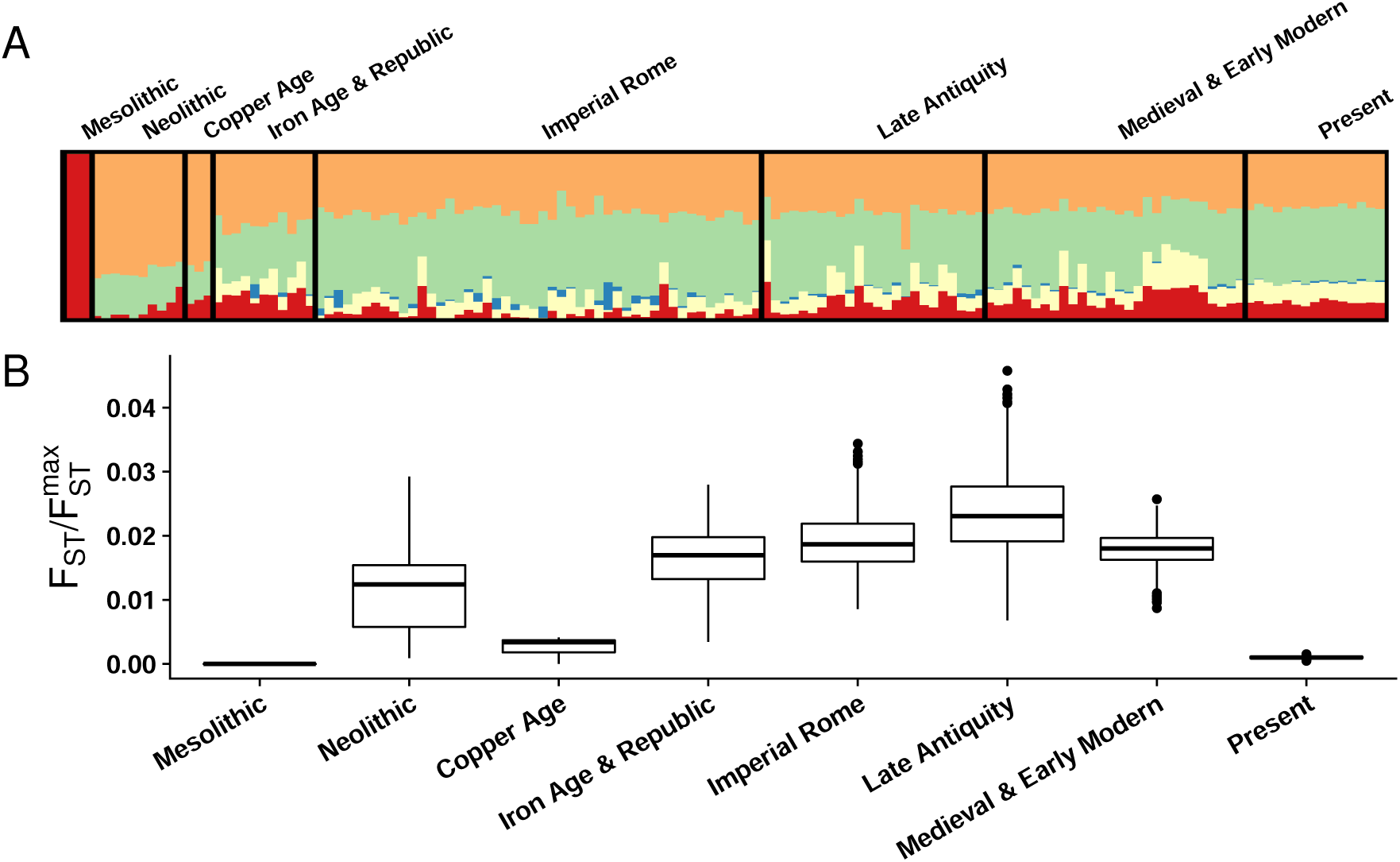
Variability of ancestry over time. **(A)** *K* = 5 Structure analysis of samples from eight time periods: Mesolithic (*n* = 3), Neolithic (*n* = 10), Copper Age (*n* = 3), Iron Age and Roman Republic (*n* = 11), Imperial Rome (*n* = 48), Late Antiquity (*n* = 24), Medieval and Early Modern (*n* = 28), and Present (*n* = 15). Adapted from Antonio *et al*. (2019). **(B)** Bootstrap distributions of the ancestry variability measure, 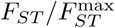, for each population (1,000 replicates).

We computed 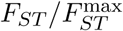 for each time period. This ratio is 0 for the Mesolithic, 0.0131 for the Neolithic, 0.0041 for the Copper Age, 0.0183 for the Iron Age and Roman Republic, 0.0192 for Imperial Rome, 0.0244 for Late Antiquity, 0.0186 for the Medieval and Early Modern period, and 0.0011 for modern individuals (Figure 6B). Pairs of bootstrap distributions of 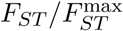 are significantly different (*p* < 2 × 10^−9^ for all 28 pairwise combinations, Wilcoxon rank sum test). The numerical results validate the claims of Antonio *et al*. (2019) of high variability during the Iron Age and Roman Republic, Imperial Rome, and Late Antiquity periods. They lend increased granularity to these claims, suggesting that ancestry variability was steadily increasing during these three periods, with a maximum achieved during Late Antiquity.

#### Different genetic loci in the same samples

The ancestry patterns identified by population structure inference methods are influenced by the choice of loci used for the analysis. When data sets possess few loci, structure is not observed, and individuals have membership coefficients close to 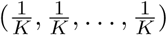; different individuals possess similar membership coefficients. As the size of the data set increases, individuals come to have different membership coefficients, with, for example, individuals from two predefined populations possessing membership primarily in two distinct clusters.

To explore patterns of ancestry variability in data sets of different size, we evaluated 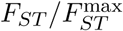 using results from a Structure analysis conducted by Algee-Hewitt *et al*. (2016). This study focused on 13 tetranucleotide loci commonly used for individual identification in forensic applications, the “Codis loci.” In a worldwide human sample, the study compared analyses with the Codis loci to analyses with a larger set of 779 non-Codis loci and to analyses with sets of 13 non-Codis tetranucleotide loci. The study claimed that the Codis loci have similar ancestry information to sets of 13 non-Codis tetranucleotide loci.

Four ancestry patterns from Algee-Hewitt *et al*. (2016), inferred from the same sample of individuals, are replotted in Figure 7. Figure 7A depicts a plot based on the Codis loci. Figure 7B plots a “null data set” designed to possess no structure. Figure 7C plots a set of 13 non-Codis tetranucleotide loci, and Figure 7D depicts a plot with 779 loci. The “null” plot shows little structure, the two plots with 13 loci show some structure, and the plot with 779 loci shows substantial structure.

**Figure 7:**
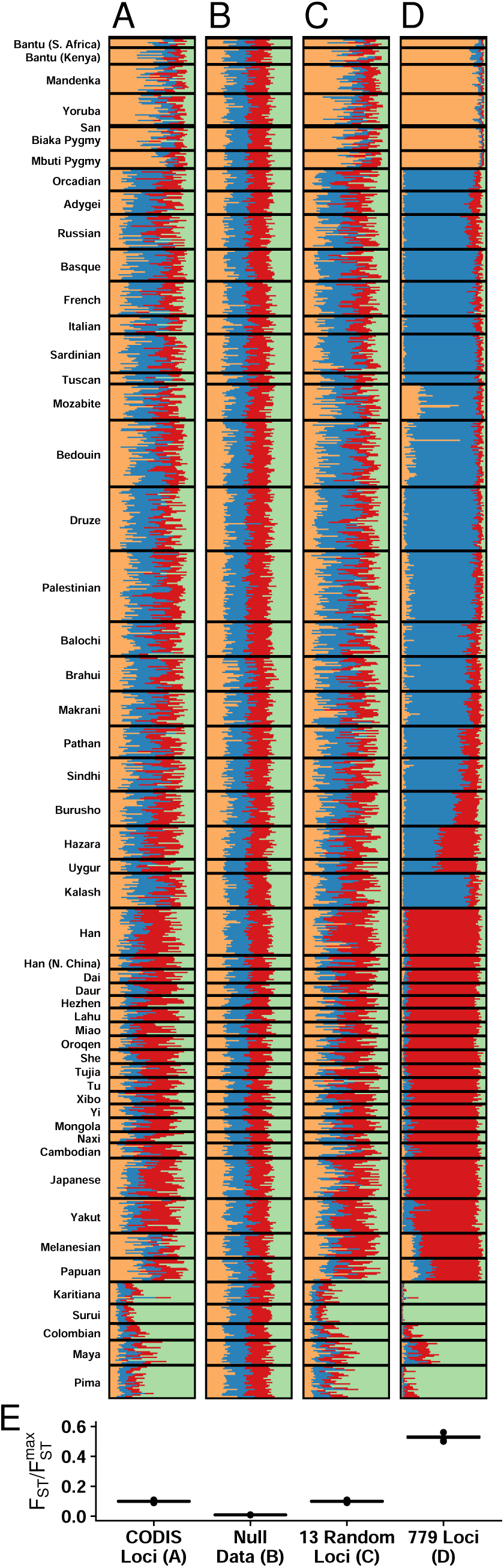
Variability of ancestry for analyses with different loci from the same samples. **(A-D)** *K* = 4 Structure analyses of four different sets of loci for a worldwide human sample. Adapted from Algee-Hewitt *et al*. (2016). **(A)** 13 Codis tetranucleotide microsatellite loci. **(B)** A simulated null data set with no population structure. **(C)** 13 non-Codis tetranucleotide microsatellite loci. **(D)** Full data set of 779 tetranucleotide loci. **(E)** Bootstrap distributions of the ancestry variability measure, 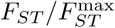, for each sample (1,000 replicates).

We computed 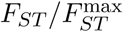 for each analysis, for each plot evaluating variability in ancestry across all individuals within the plot. The ratio is lowest for the null data set, with a value of 0.009. It is 0.100 for both the Codis loci and for the 13 non-Codis loci. The ratio is substantially higher for the full 779 loci, with a value of 0.529. Five of the six pairs of bootstrap distributions of 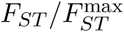 are significantly different (*p* < 2×10^−16^, Wilcoxon rank sum test), the exception being that the two plots with 13 loci, Codis and non-Codis, do not show a significant difference (*p* = 0.56). The pattern of 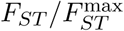 values, with the smallest value for Figure 7B, intermediate values for Figure 7A and Figure 7C, and largest value for Figure 7D, captures increasing ancestry variability as the analyses move from a largely unstructured plot (Figure 7B) to partially unstructured plots (Figure 7A and 7C) to a substantially structured plot (Figure 7D). The lack of a significant difference in 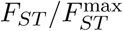 between the plot for the Codis loci and the plot for equally many non-CODIS loci supports the claim of Algee-Hewitt *et al*. (2016) that the Codis loci contain comparable information about ancestry to other sets of loci with the same size.

## Discussion

We have introduced a measure for quantifying variability across vectors of individual membership coefficients, as produced by population structure inference programs such as Structure and Admixture. Our measure is based on a mathematical analogy with the population differentiation statistic *F*_*ST*_. Whereas *F*_*ST*_ traditionally measures variability in allele frequency vectors among populations, we have used *F*_*ST*_ to measure variability in membership coefficient vectors among individuals. Because the upper bound of *F*_*ST*_ as a function of the frequency of the most frequent allele is usually less than 1, we have employed a normalized version of this statistic, 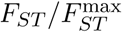, which ranges in [0, 1] for all matrices of membership coefficients and can thus be used to compare ancestry variability among different matrices.

Through both simulation and an analytical calculation under a Dirichlet distribution for membership coefficient vectors, we demonstrated that the expected value of 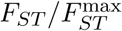 increases with variance of membership coefficients across individuals (Figure 3 and 4); indeed, it scales approximately linearly with the parametric variance in a model with *K* = 2 ancestral clusters (eq. 2). This result supports the use of 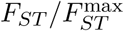 as a measure of variability in ancestry across individuals.

We also demonstrated the use of the measure in data sets exemplifying three scenarios in which ancestry variability is of particular interest. In a comparison of ancestry measured in admixed and non-admixed populations by Verdu *et al*. (2017), we found that the recently admixed Cape Verdean population exhibited greater variability in ancestry, as measured by 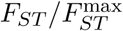, than did non-admixed populations (Figure 5). In a comparison of ancestries measured in different time periods in the same location, we provided quantitative support for a claim of Antonio *et al*. (2019) that certain eras in ancient Rome possessed more variable ancestry than others (Figure 6). Finally, in a comparison of different sets of loci studied in the same individuals, we found quantitative support both for the observation of Algee-Hewitt *et al*. (2016) that ancestry variability across individuals was similar for two different sets of 13 loci, and for an increase in ancestry variability in high-resolution data compared to data of lower resolution. In all three cases, our analyses provided quantitative support for claims previously argued primarily by qualitative observation.

Because the measure depends on *Q* matrices, limitations of the methods used to generate the *Q* matrices extend to our measure. For example, if individuals were mislabeled prior to analysis with methods such as Structure or Admixture, then our measure would be affected. Further, *Q* matrices generated by Structure and Admixture do not contain information about the magnitude of the difference between ancestral clusters; our measure only captures variation in ancestry with respect to the clusters that such programs infer.

The new measure, which we have implemented in the *R* package FSTruct, contributes to a body of methods for quantitative analysis of inferred membership coefficients. This collection of methods includes computations useful for analyzing the level of support observed for different numbers of clusters *K* (Evanno *et al*., 2005) and methods of aligning the clustering solutions observed in replicate analyses (Jakobsson and Rosenberg, 2007; Kopelman *et al*., 2015; Behr *et al*., 2016), as well as software for graphical display (Rosenberg, 2004) and for managing files associated with the analysis (Earl and VonHoldt, 2012).

A number of other studies have considered related but distinct problems in assessing variability of ancestry based on membership fractions. Rosenberg *et al*. (2005) described a “clusteredness” statistic that measures the extent to which individuals are placed into single clusters rather than across multiple clusters. This statistic is maximal if each individual possesses a permutation of the membership vector (1, 0, …, 0) and minimal if all individuals possess membership vector 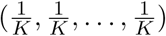. Kerminen *et al*. (2021) evaluated the Shannon entropy applied to individual-level membership vectors, assessing variation in time in the Shannon entropy for study participants with different birth years. Whereas both the clusteredness statistic of Rosenberg *et al*. (2005) and the Shannon entropy statistic of Kerminen *et al*. (2021) consider variability of the ancestry coefficients of single individuals, our 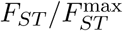 measure examines variability of ancestry coefficient vectors *across* individuals. Thus, for example, comparing individuals in corresponding matrices in Figure 4A and 4D, clusteredness increases (and Shannon entropy decreases) as the membership of the highest-membership cluster increases from Figure 4A to Figure 4D. However, 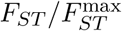, measuring variability *across* individuals, is similar in corresponding matrices in the two panels, reflecting the visual similarity between panels of the inter-individual patterns.

We note that in addition to analyzing the *Q*-matrices produced by population structure inference programs such as Structure and Admixture, FSTruct can quantify variability in any matrix whose rows sum to 1. Applications are potentially numerous. For example, single-cell sequencing technologies have enabled the identification and quantification of cell populations within tissues, revealing different patterns of variation, with some tissues containing few cell populations, while others are more diverse (Wang *et al*., 2019). Our method enables comparisons of the variability of within-tissue cell populations, where tissues are analogous to individuals and cell populations are analogous to cluster memberships. Our method could also be applied to quantify variability among individuals of features such as mutational signatures, where the proportion of mutations belonging to a mutational type is analogous to a cluster membership (Alexandrov *et al*., 2013; Rahbari *et al*., 2016).

## Appendix

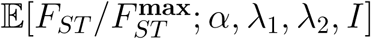

In this appendix, we evaluate the approximate expected value of 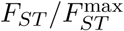 calculated for a sample of *I* individuals (*i* = 1, 2, …, *I*), each with *K* = 2 membership coefficients 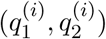 drawn independently from the Dirichlet distribution Dir(*αλ*_1_, *αλ*_2_). We use the notation 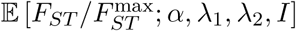 or simply 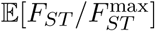 to denote this expectation.

### Overview

To obtain the expectation 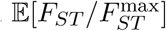, we first sample *I* independent and identically distributed Dir(*αλ*_1_, *αλ*_2_) random variables 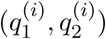, where 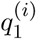 is the membership coefficient of individual *i* in cluster 1, 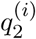 is the membership coefficient of individual *i* in cluster 2, and 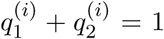. We assume that the sample size *I* is large.

We assume without loss of generality that the parametric mean membership coefficient for cluster 1 is at least as large as that for cluster 2; that is, *λ*_1_ ≥ *λ*_2_. As *I* → ∞, by the strong law of large numbers (Serfling, 1980, section 1.8), the sample mean membership coefficient for cluster 1, 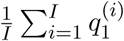, converges almost surely to the parametric mean *λ*_1_, and the sample mean membership coefficient for cluster 2, 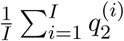, converges almost surely to the parametric mean *λ*_2_. Hence, for large *I*, the probability approaches 1 that 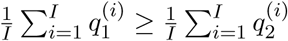. As a result, because we consider large *I*, we assume that the cluster with the greater *parametric* mean membership coefficient, cluster 1, also has the greater *sample* mean membership coefficient. We denote this sample mean, the mean membership of cluster 1 in a simulated population, by 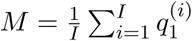. By definition, 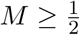. As stated above, 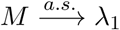 as *I* → ∞.

The quantity whose expectation we wish to evaluate under the model, 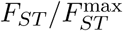, is a function of the sampled membership coefficients, 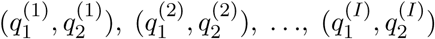. We let 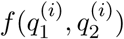 represent the Dirichlet probability density for 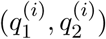; because we are considering vectors with two components, the Dirichlet reduces to a Beta distribution (Kotz *et al*., 2000, p. 487),

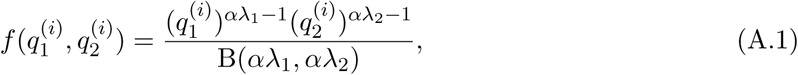

where

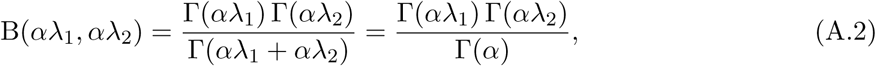

and Γ is the gamma function.

The 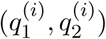 are independent and identically distributed. Hence, the expectation is

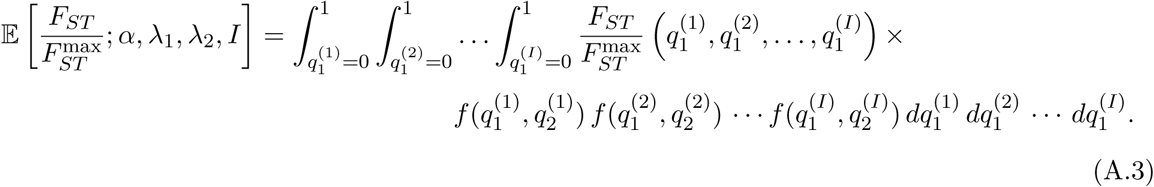

With this expression in hand, we proceed by writing the expression for 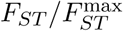 in terms of the membership coefficients 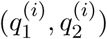 and the sample size *I*. We then compute the integral, making use of the Dirichlet parameters *α, λ*_1_, and *λ*_2_.

### Approximating 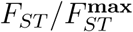 under the Dirichlet model

The value of 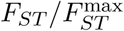 calculated for a population of *I* individuals with membership coefficients 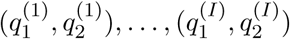, can be written using eqs. 3 and 5 of Alcala and Rosenberg (2017),

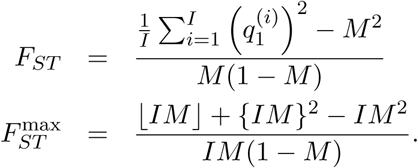

We obtain

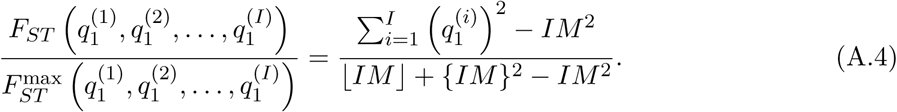

Recall that 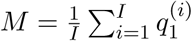 is the sample mean membership of the most prevalent ancestral cluster, assuming that the cluster with the greater parametric mean membership is also the cluster with the greater sample mean membership.

We now make an approximation to the denominator of eq. A.4. Because ⌊*IM* ⌋ = *IM* − {*IM*}, ⌊*IM* ⌋+{*IM*}^2^ = *IM* −({*IM*} −{*IM*}^2^) = *IM* − *δ* where the error term *δ* = {*IM*}(1 −{*IM*}) lies in 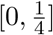, taking its maximal value of 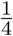 when 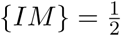. For large sample size *I*, because 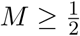 and 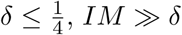, so that ⌊*IM* ⌋ + {*IM*}^2^ ≈ *IM*. Thus, we substitute *IM* in place of ⌊*IM* ⌋ + {*IM*}^2^ in eq. A.4, obtaining

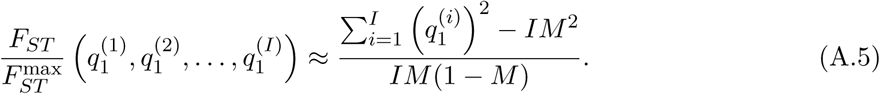

This assumption is equivalent to setting 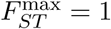.

To find an approximation for 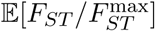, it is convenient to make a further approximation in eq. A.5, substituting *M* with *λ*_1_. We justify this substitution by proving that as *I* → ∞,

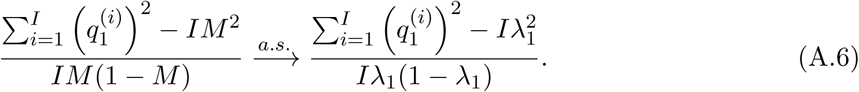

Subtracting the right-hand side from the left-hand side, proving eq. A.6 is equivalent to proving

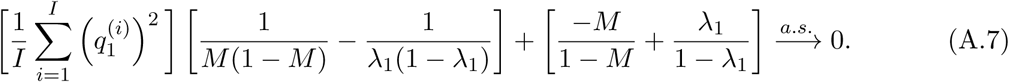

If 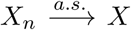 and 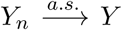, then 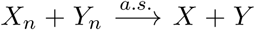 (Grimmett and Stirzaker 2001a, p. 336, exercise 2; Grimmett and Stirzaker 2001b, p. 354, exercise 2), so the sum of two terms that converge almost surely to 0 also converges almost surely to 0. Hence, it suffices to separately prove almost sure convergence to 0 of the two terms summed in eq. A.7.

For the right-hand term, we use the continuous mapping theorem, which states that for a continuous function *g* and a random vector *X*_*n*_, if 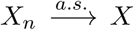 then 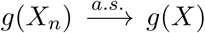 (VAN DER Vaart, 1998, p. 7, Theorem 2.3). We consider the continuous function *g*(*x*) = −*x/*(1 − *x*) + *λ*_1_*/*(1 − *λ*_1_) and recall that as *I* → ∞, 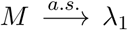. It follows that as *I* → ∞, and 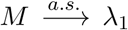, 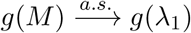; that is, 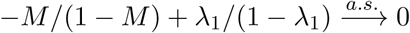.

For the left-hand term, the factor 1*/*[*M*(1 − *M*)] − 1*/*[*λ*_1_(1 − *λ*_1_)] converges almost surely to 0 by the continuous mapping theorem with *g*(*x*) = 1*/*[*x*(1 − *x*)] − 1*/*[*λ*_1_(1 − *λ*_1_)]. By the strong law of numbers, 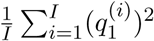 converges almost surely to 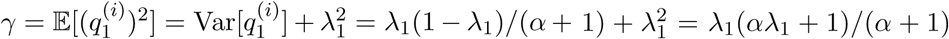 (Serfling, 1980, Theorem B). If 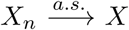 and 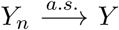, then 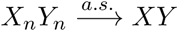 (Grimmett and Stirzaker 2001a, p. 336, exercise 2; Grimmett and Stirzaker 2001b, p. 354, exercise 2), so that the left-hand term of eq. A.7 converges almost surely to *γ* × 0 = 0.

### Evaluating 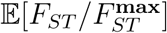 under the Dirichlet model

Inserting our expression for 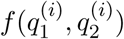 from eq. A.1 and our expression for 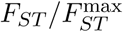 from eq. A.6 into eq. A.3 allows us to write an approximate expression for the expectation of 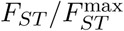 given the parameters of the Dirichlet distribution and the sample size *I*:

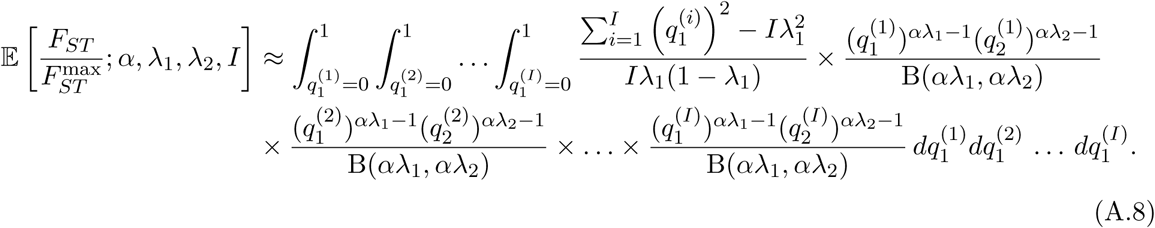

Examining the quantity 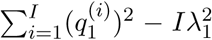, we observe that eq. A.8 can be decomposed as a sum of *I* + 1 terms, one for each of the 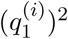 terms, and one for the 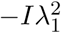 term. Assign the first *I* of these separate terms the labels *L*_1_, *L*_2_, …, *L*_*I*_ and the 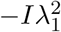 term the label *L*_∗_, so that 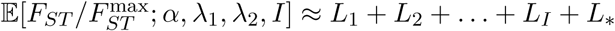.

We begin by evaluating the term *L*_∗_, which can be written

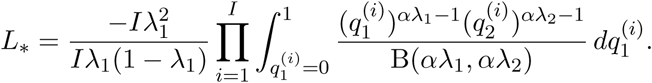

Recalling that 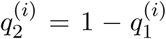, we observe that the integrand 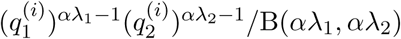 is simply the Beta probability density function, which integrates to one. Hence, the product evaluates to 1 and *L*_∗_ simply equals a constant:

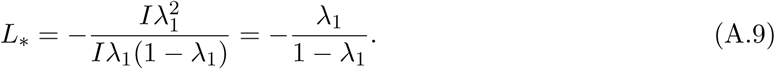

We next evaluate the *L*_*i*_ terms. For each *i* in 1, 2, …, *I*,

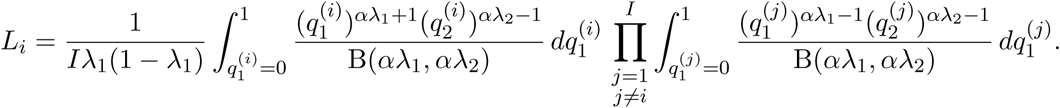

As was the case for *L*_∗_, the integrand of the integral inside the product is the Beta probablility density function, so the product evaluates to one. Thus,

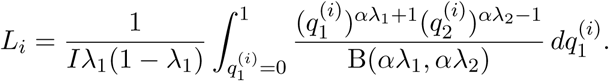

The remaining integral can be evaluated by noting that 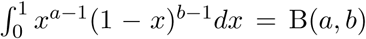. We employ this identity to simplify *L*_*i*_, obtaining

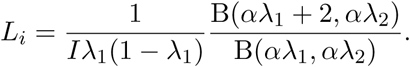

By eq. A.2 and the property of gamma functions Γ(*z* + 1) = *z*Γ(*z*), this expression simplifies to

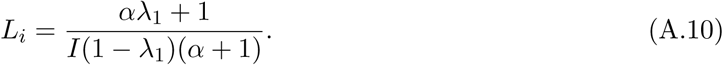

We now combine eqs. A.9 and A.10 to complete the calculation in eq. A.8, noting that *L*_*i*_ does not depend on *i*, so that each *L*_*i*_ follows eq. A.10.

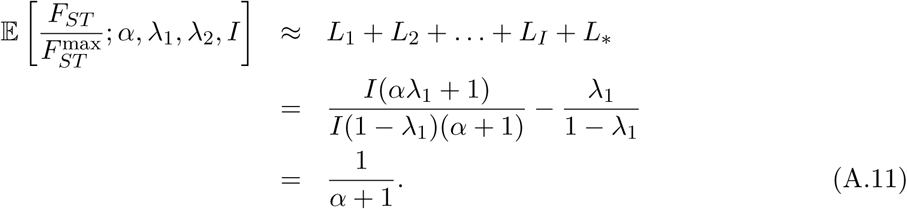

## Acknowledgments

We acknowledge support from NIH grant R01 HG005855. MLM acknowledges support from a National Science Foundation Graduate Research Fellowship and the Anne T. and Robert M. Bass Stanford Graduate Fellowship. We thank P. Verdu, M. Antonio, and M. Edge for assistance with data sets from their studies, H. Moots and J. Pritchard for helpful comments on the method, and D. Cotter and J. Mooney for suggestions for the software. Where authors are identified as personnel of the International Agency for Research on Cancer/World Health Organization, the authors alone are responsible for the views expressed in this article and they do not necessarily represent the decisions, policy, or views of the International Agency for Research on Cancer/World Health Organization.

## Data accessibility

The FSTruct R package is available for download from github.com/MaikeMorrison/FSTruct. The introductory vignette is linked from the package README file and provides a guide to use of the package.

## Author contributions

MLM, NA, and NAR designed the study and performed the theoretical analysis. MLM conducted the simulations, analyzed the data, and wrote the software. NAR supervised the study. All authors wrote the manuscript.

## Notes

https://github.com/MaikeMorrison/FSTruct

### Competing Interest Statement

The authors have declared no competing interest.

